# Crispr2vec: Machine Learning Model Predicts Off-Target Cuts of CRISPR systems

**DOI:** 10.1101/2020.10.28.359885

**Authors:** Tara Basu Trivedi, Ron Boger, Govinda M. Kamath, Georgios Evangelopoulos, Jamie Cate, Jennifer Doudna, Jack Hidary

**Affiliations:** Alphabet (Google) X, Mountain View, CA, 94043, USA; Department of Molecular and Cell Biology, University of California, Berkeley, Berkeley, CA, 94720, USA; Department of Chemistry, University of California, Berkeley, Berkeley, CA, 94720, USA; Innovative Genomics Institute, University of California, Berkeley, Berkeley, CA, 94720, USA

## Abstract

Clustered Regularly Interspaced Short Palindromic Repeats (CRISPR)/CRISPR-Cas systems have revolutionized gene editing, with applications in therapeutics, diagnostics, agriculture, and developing disease models. However, CRISPR-Cas suffers from off-target effects — unintended genetic modifications in the genome that arise from its use. In this work, we present crispr2vec: a deep metric learning approach for embedding CRISPR single guide RNA (sgRNA) sequences and predicting off-target cuts. Given a fixed target sequence, we show that our learned embedding yields a faithful representation of potential off-targets. We present a new triplet sampling strategy specifically for CRISPR sequences that improves the quality of our embedding. We show the resulting embedding generalizes across different off-target cut detection assays. Finally, we demonstrate the superiority of our deep metric learning method in its ability to predict off-target cuts compared to previous literature in cross fold validation across different datasets for both seen and unseen sgRNAs.

## 2 Introduction

Since their description as a genome editing technology, Clustered Regularly Interspaced Short Palindromic Repeats (CRISPR)-CRISPR-associated (Cas) systems [24, 16] have enabled more efficient and effective genome engineering than was previously possible [34, 11, 41, 22]. The CRISPR-Cas system’s ability to target specific sequences in DNA and RNA in organisms has led to its widespread use, with applications of CRISPR-Cas enzymes now including diagnostics [5, 25], therapeutics [50, 43, 38], agriculture [20], cell models [14, 7], and more.

Fundamentally, this technology relies on RNA-guided enzymes that recognize and cut DNA or RNA molecules whose sequence matches the sequence of a ~ 20-nucleotide segment within a guide RNA. CRISPR-Cas9, the first such system harnessed for genome editing, cuts DNA at positions defined by the guide sequence in single-guide RNAs (sgRNAs) that were engineered to create a simple two-component system for use in cells. Cas9 introduces a blunt double-stranded cut in DNA within the sequence complementary to the guide RNA and adjacent to a short protospacer-adjacent motif (PAM) sequence. For the commonly used Cas protein SpCas9, the PAM sequence is NGG (see Figure 1a for more details). There are now a range of CRISPR systems making use of different Cas proteins, all of which use guide RNAs [30].

**Figure 1:**
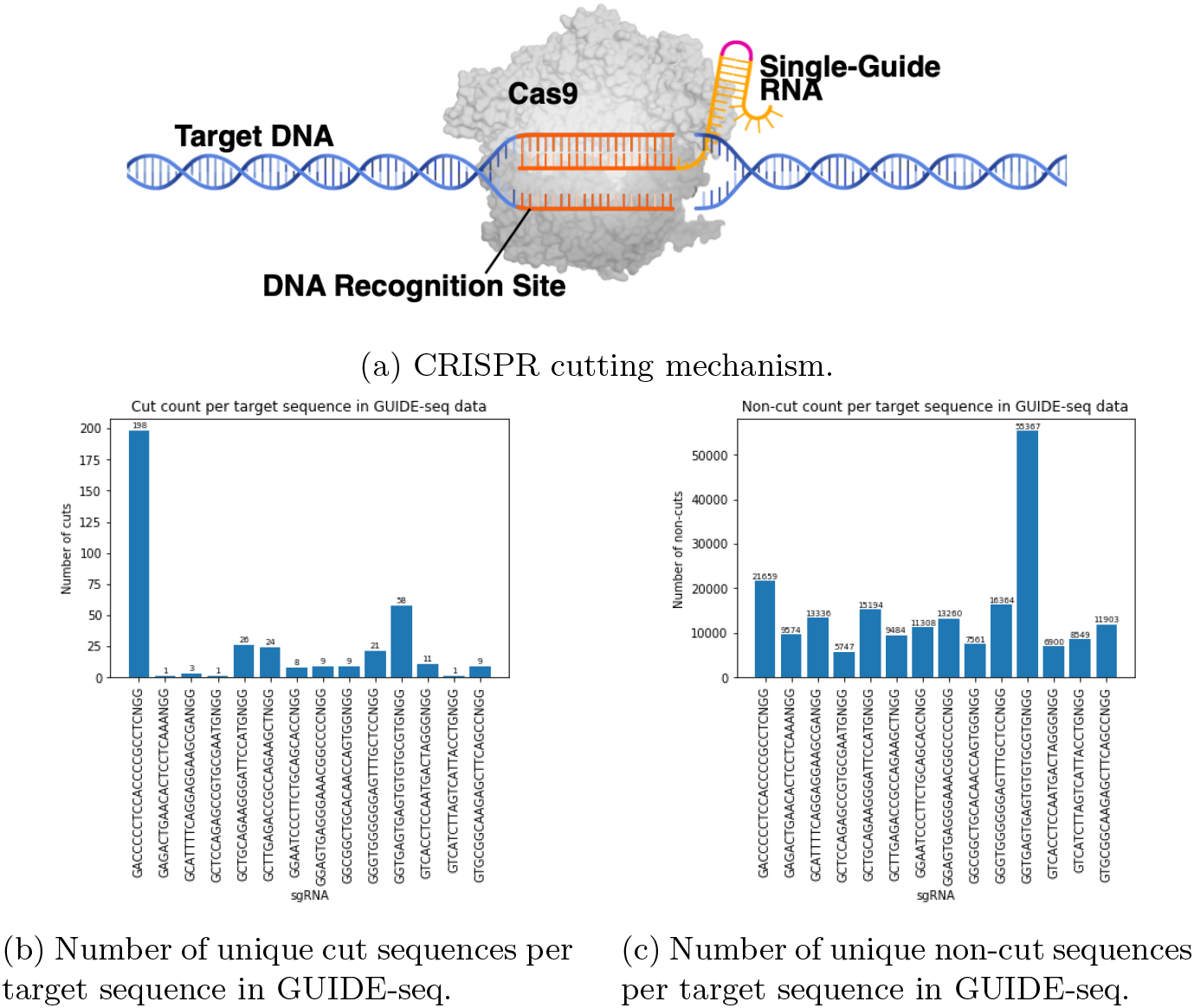
CRISPR cut sites.

Despite its widespread utility for genome editing applications, CRISPR-Cas9 presents challenges in its sensitivity and specificity. The Cas9 protein does not always cut all DNA sequences in a cell that are complementary to the sgRNA guide sequence. Additionally, the Cas9 protein sometimes cuts positions in the genome that are not fully complementary to the sgRNA sequence [10]; we refer to these cuts as off-target effects. These off-target effects that can result from the use of CRISPR-Cas9 can lead to unintended gene alterations. Thus, being able to predict and control whether a given sgRNA will produce off-target effects is paramount to any clinical use of CRISPR-Cas9.

There are many *in-vitro* assays to profile off-target DNA cuts. The most popular ones include GUIDE-seq [48, 29, 6], which is based on integration of oligonucleotides into double strand breaks detected by sequencing; Digenome-seq [21, 27], based on in-vitro nuclease-digested whole-genome sequencing; high-throughput genome-wide translocation sequencing or HTGTS [18]; BLESS-seq [40, 45], based on direct in situ break labeling; integration-deficient lentiviral vectors or IDLV [49]; and CIRCLE-seq [47], which provides a highly sensitive in-vitro biochemical assay that does not require a reference genome sequence and can be used to identify off-target mutations associated with cell type-specific single nucleotide polymorphisms.

Given the combinatorial number of possible CRISPR sequences and off-target effects, the space of sequences is too large to search. Being able to predict if an sgRNA will cut other areas of the genome allows a biologist to quickly eliminate sgRNA sequences and increases confidence around the clinical use of CRISPR. This presents major challenges, since there is not much data and the relevant features are unclear. Our work thus focuses on *in-silico* profiling, leveraging datasets from several of the aforementioned *in-vitro* assays.

## 3 Prior State of the Art

A number of solutions have been presented for the off-target prediction problem, including Hsu-Zhang [23], Cas-OFFinder [4], CHOPCHOP [35], CFD [15], CRISPOR [21], CRISTA [1], Elevation [32], DeepCrispr [19], and DeepSpCas9 [28]. Each vary in their chosen searching and scoring algorithms.

- Hsu-Zhang scores [23] predict cleavage of off-target sequences based on experimental cleavage affinities of mismatches at each position along the sgRNA and the DNA sequences.
- Cas-OFFinder [4] scalably identifies potential off-target sites by searching the genome for similar sequences, allowing some number of mismatches in sequence alignments.
- CHOPCHOP scores [35] offer predictions of off-target binding of sgRNAs using efficient sequence alignment algorithms.
- CFD scores [15], or “cutting frequency determination” scores, estimate the likelihood of off-target activity by considering experimentally observed percentage activity between mismatches on sgRNA sequences and DNA target sequences.
- For each target sequence, CRISPOR [21] calculates a specificity score and two efficiency scores. The specificity score captures the likelihood of off-target cleavage elsewhere in the genome resulting from the target sequence’s associated sgRNA sequences; they use the Hsu-Zhang score [23]. The efficiency score captures the likelihood of the target sequence being cleaved by its associated sgRNA sequences; they use CFD [15] and CRISPRscan [36].
- CRISTA [1] estimates the propensity for genomic sites to be cleaved by sgRNA using a gradient-boosted random forest model. In contrast to earlier works, CRISTA [1] considers additional attributes (beyond sequence alignment) such as spatial structure and rigidity of the site.
- Elevation [32] predicts the “goodness” of sgRNA sequences. The approach uses a gradient-boosted random forest model to identify target sequences’ off-target pairs and then combines the scores of all off-targets within a Hamming distance to generate a specificity score.

The last few years have witnessed the rising impact of deep learning, due to increased computational power, larger dataset sizes, and the ability to model arbitrary functions. In particular, deep learning approaches have proven superior to previous machine learning methods in fields such as computer vision [31], natural language processing [12], music generation [13], and speech recognition [37].

Deep learning has also recently outperformed statistical and classical machine learning approaches in computational biology, such as predicting the sequence specificities of nucleic acid binding proteins [3, 51], improving variant calling in next generation sequencing [39], understanding effects of non-coding sequences [53, 52], predicting gene expression from histone modification [44], and learning functional activity of DNA sequences [26]. We refer the reader to [54, 17] for a more comprehensive overview of deep learning in genomics. Deep learning has also been applied to better understand the mechanisms of CRISPR.

- Similar to Elevation [32], DeepCrispr [19] captures the “goodness” of an sgRNA sequence and its effectiveness at knocking out genes. DeepCrispr [19] fully automates the identification of sequence features and epigenetic features, using first an auto-encoder to learn sequence embeddings and then feeding the embeddings into a 9-layer convolutional neural network.
- DeepSpCas9 [28] predicts SpCas9 protein activity; it claims to be more generalizable than DeepCrispr [19]. DeepSpCas9 [28] compiled a large lentiviral library, observed sgRNA-directed SpCas9 cleavage of the cell library, and used that data to train a deep learning-based regression model to predict SpCas9 activity.

## 4 Methods

In our analysis of previous literature, we noticed several shortcomings. Our methods overcome these and depart from the prior state of the art as follows:

- A number of the previous methods require hand-crafted feature engineering. We learn a distance between sequences parameterized by a deep neural network, which enables crispr2vec to learn a more robust representation of the sequences themselves without the need for hand-crafted features.
- A key methodological issue with prior approaches is that they frame off-target detection as a simple classification problem, without consideration of the target itself. In contrast, crispr2vec considers the target sequence itself and determines inherent cutting dynamics *relative* to the target.
- Instead of unifying datasets from different off-target detection assays, we treat these approaches independently and show that our model is flexible enough to generalize across them. This is due to crispr2vec learning a distance from the intended CRISPR target rather than a sequence classification.

### 4.1 Datasets

For this paper, we first restricted ourselves to GUIDE-seq [48, 29, 6] data for target sequences and off-target cut sequences. We only considered unique target-off-target sequence pairs - that is, the only features we considered come from the sequences themselves; not, for example, their location in the genome. We then used Cas-OFFinder [4] to find sequences in the genome that are within a Hamming distance of 6 from the target sequence under consideration. We filtered out target-off-target sequence pairs from Cas-OFFinder that overlapped with pairs in the GUIDE-seq data set to get a final list of non-cut off-target sequences. This left us with 395 unique off-target cut sequences (Figure 1b) and 206,206 unique off-target non-cut sequences (Figure 1c) across 14 target sequences. In addition, for the purposes of training and testing, we filtered out target sequences that were associated with too few unique off-target cuts.

Later, we pulled cleavage data from the union of GUIDE-seq and CIRCLE-seq [47] data in order to examine the ability of our method to generalize to other off-target cut detection assays. We chose the CIRCLE-seq assay in particular due to the large number of off-targets detected (which makes it more amenable to our deep learning approach), as well as its greater sensitivity than GUIDE-seq (since off-targets found in GUIDE-seq are effectively a subset of those in CIRCLE-seq). Again, we only considered unique target-off-target sequence pairs. Our non-cut target-off-target sequence pairs came from filtered Cas-OFFinder data that did not overlap with pairs in the GUIDE-seq and CIRCLE-seq data. This left us with 5,684 unique off-target cut sequences (Figure S1a) and 217,855 unique off-target non-cut sequences (Figure S1b) across 18 target sequences. In addition, for the purposes of training and testing, we filtered out target sequences that were associated with too few unique off-target cuts.

Each DNA sequence was one-hot encoded into an array of integers that could be input into our model. (See Figure 2a for an example encoding).

**Figure 2:**
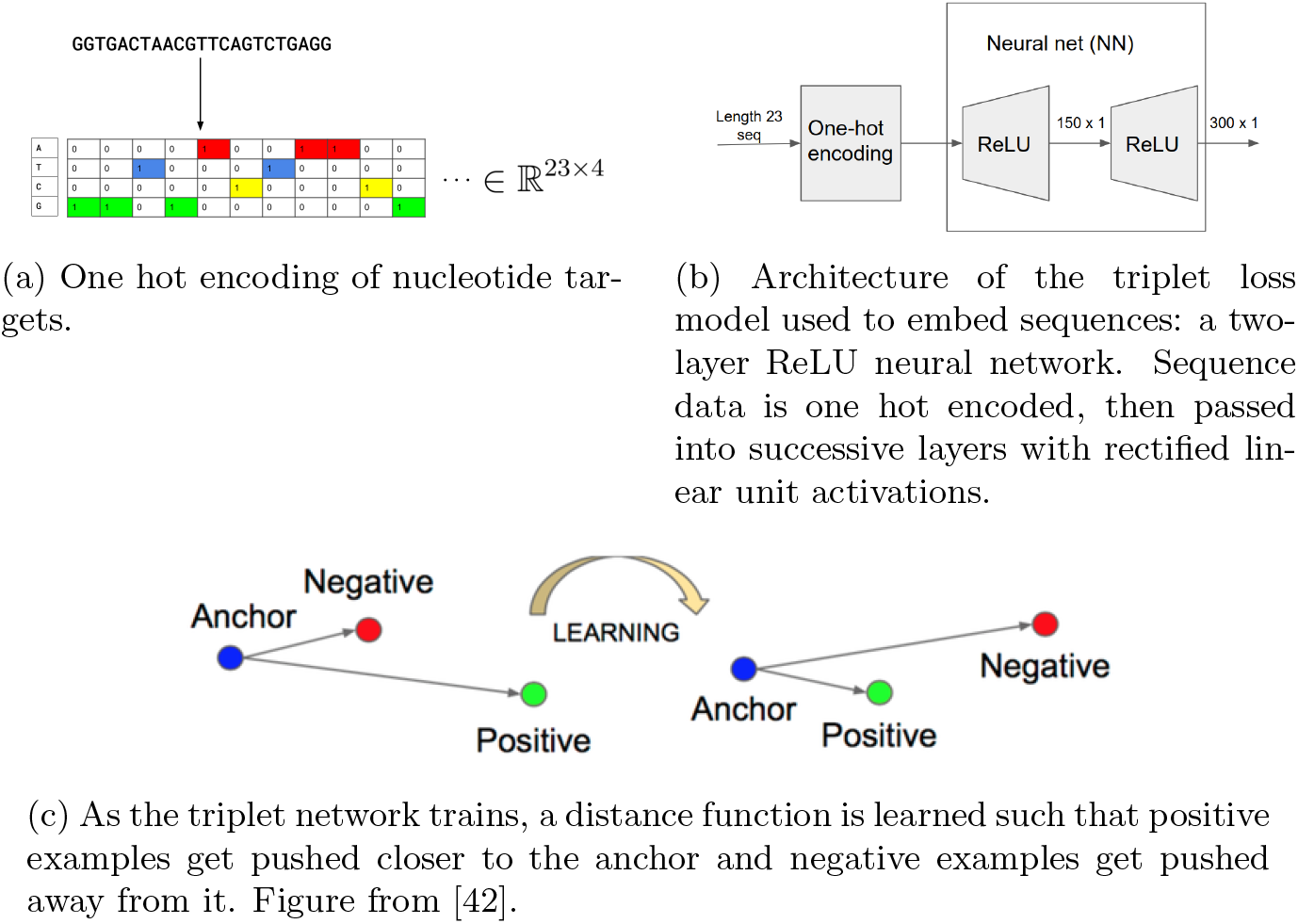
Summary of our data processing platform.

For each target sequence, we generated 4 one-hot encodings; one for each A/T/G/C sequence variation. For each off-target sequence, we generated a single one-hot encoding.

### 4.2 Model

Recent literature for CRISPR off-target predictions focuses on directly classifying off-target sequences for a particular sgRNA. The contrasts with another approach called metric learning, which is the task of learning a distance function between objects in data. We hypothesized that metric learning is a more natural way to frame the problem for the following reasons:

- Off-target sites that are cut are presumably “close” to the desired on-target cut site with an appropriate distance metric.
- The classification regime does not naturally take into consideration the desired on-target sequence.
- Metric learning easily admits the use of deep learning to learn a distance between CRISPR sequences.
- Metric learning models are famous few-shot learners, i.e. they still achieve state of the art performance where each category is represented by only a few examples.

We use triplet loss, a metric learning approach that enables use of a neural network to generate an embedding that produces distances between sequences (Figure 2b). This was first introduced in the context of face recognition [42]. Deep metric learning approaches have shown wide applicability, ranging from face re-identification [9, 8] to computational biology [2] and perform strongly in recognition and one-shot learning tasks. At each step in the training process, triplets of data are passed into the model. In our case, the triplets are the intended cut site (termed the “anchor”), off-target cut site identified by GUIDE-seq and CIRCLE-seq (“positive example”), and off-target non-cut site identified by Cas-OFFinder [4] (“negative example”). The goal of the training step is to minimize the learned distances between the intended cut sites and the off-target cut sites, while maximizing the learned distances between the intended cut sites and the off-target non-cut sites. (See Supplementary methods section 1.2 for more information).

Given the loss function for which we are optimizing, choosing which triplets to pass into the network can be the difference between a highly expressive model and a degenerate one. Much of the existing literature on this topic focuses on methods for semi-hard triplet mining, both in offline (precomputed) and online (estimated with each batch) settings. We use a novel triplet sampling method that we term **smart mining**, which samples sequences in accordance to their biological feasibility in the CRISPR off-target prediction problem. (See Supplementary methods section 1.2.1 for more information).

We performed cross validation to assess the performance of our model relative to state of the art baselines. (See Supplementary methods sections 1.2.2 and 1.2.3) for more information).

## 5 Results

See Table S1 for a summary of all crispr2vec results (including AUC-ROC values, precision values, recall values, and F1 scores).

### 5.1 Testing on Seen sgRNAs

We started by considering only GUIDE-seq for positive examples. With a margin hyperparameter *α* of 1.0, the loss function quickly converged (Figure 4a). For each fold, we examined the calculated distances between anchors to positive examples and anchors to negative examples in the held-out test set (Figure 4b). This was a useful indicator for how well the model had learned embeddings and, by extension, distances between sequences. We performed an area under the receiver operating characteristic curve (AUC-ROC) analysis on the cross validation experiment. AUC-ROC is a performance metric that captures how much the model is able to distinguish between classes. The higher the AUC-ROC value, the better the model is at classifying cut sequences as cut sequences and non-cut sequences as non-cut sequences (note that random performance corresponds to an AUC-ROC of 0.50). The average AUC-ROC value was 0.96, with a standard deviation of 0.012 (Figure 3a, Table S1). We compared crispr2vec to the aforementioned baseline models and classical machine learning methods. We do not compare to methods such as deepCRISPR [19], DeepSpCas9 [28], and Elevation [32]. We note that while these methods would theoretically serve as good benchmarks for our method, they both use different data sources (for instance, integration of epigenetics) and the trained models these papers provide online are trained on guide-RNAs contained in the test set for our method in each scenario.

**Figure 3:**
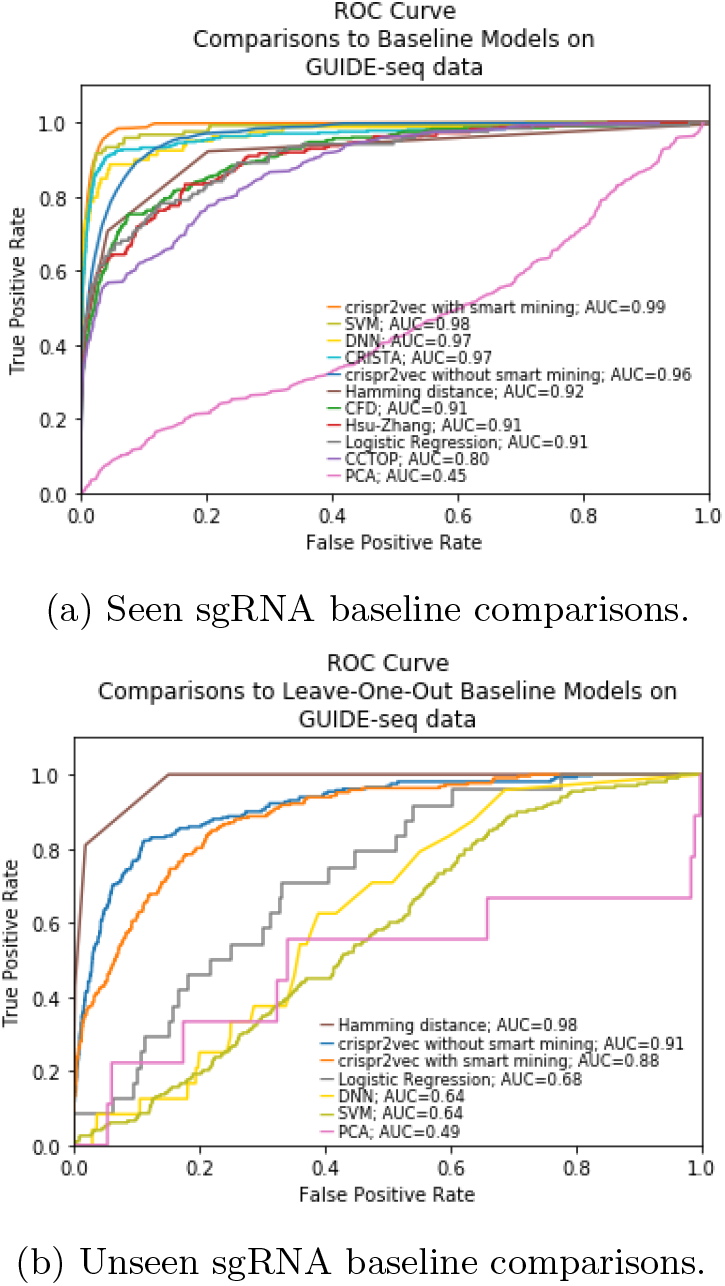
Comparison of AUC-ROC values for crispr2vec trained on GUIDE-seq data versus baseline models.

**Figure 4:**
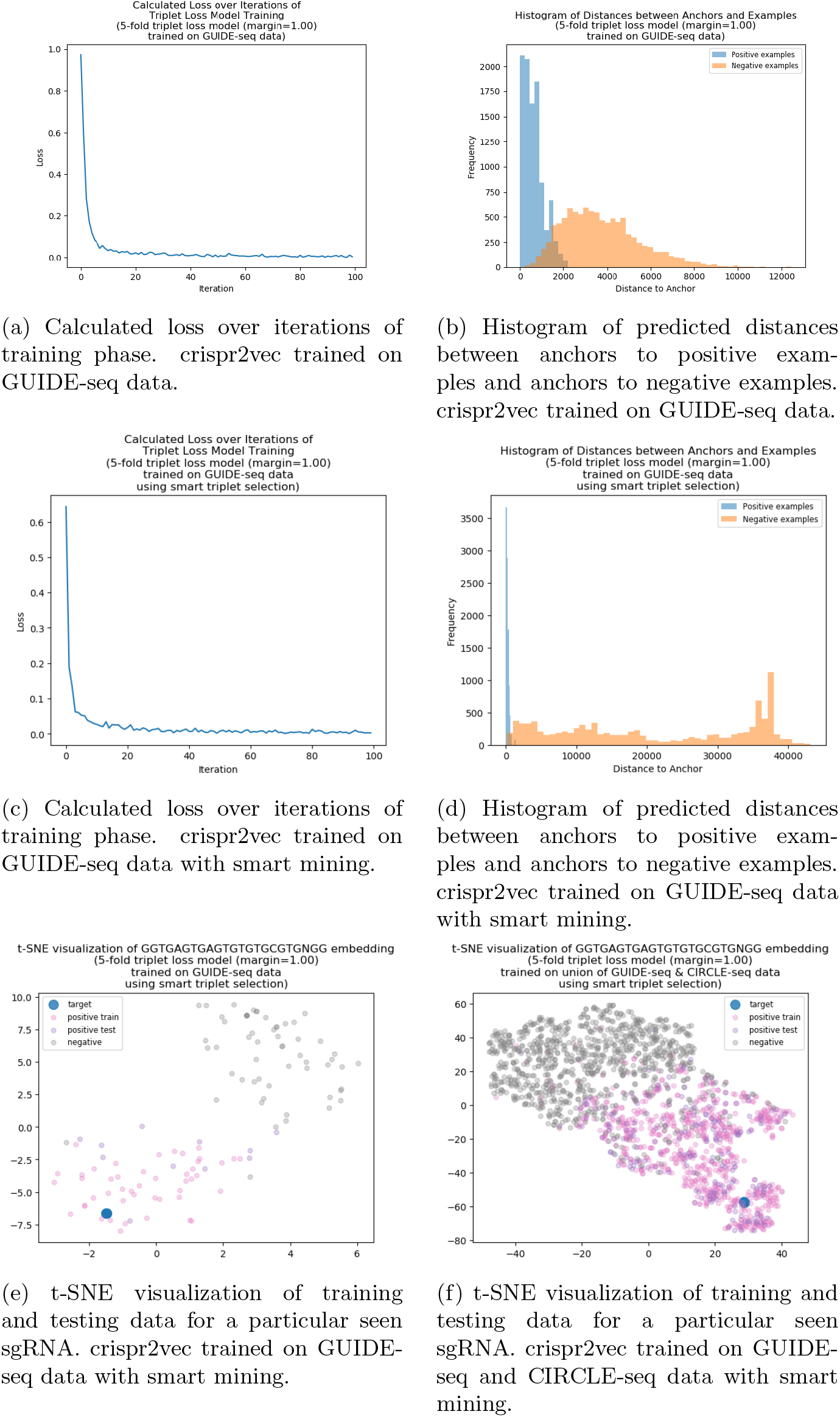
5-fold cross validation of crispr2vec with and without smart mining.

#### 5.1.1 Smart Mining

Next, we added smart mining (refer to Supplementary section 1.2.1) to see if it enhanced the model’s ability to classify sequences. With a margin hyperparameter *α* of 1.0, the loss function quickly converged (Figure 4c). For each fold, we examined the calculated distances between anchors to positive examples and anchors to negative examples in the held-out test set. We noticed that the spread of anchor-to-negative-example distances was much larger for the smart mining model (Figure 4d) than for the base triplet loss model (Figure 4b). In addition, the difference between the mean of the anchor-to-positive-example distances and the mean of the anchor-to-negative-example distances for the smart mining model (Figure 4d) was much larger than that of the base triplet loss model (Figure 4b). The average AUC-ROC value was 0.99, with a standard deviation of 0.0041 (Table S1). When testing on seen target sequences, the triplet loss model that employed smart mining had a higher AUC-ROC than did the base triplet loss model (Figure 3a).

We employed the t-distributed stochastic neighbor embedding (t-SNE) [33] algorithm to visualize the embeddings learned by crispr2vec. We visualized the embeddings of training and testing data for target sequences (Figure 4e, 4f).

When comparing with baseline models, we found that crispr2vec with smart mining had the highest AUC (Figure 3a).

### 5.2 Testing on Unseen sgRNAs

We wanted to validate crispr2vec on unseen guide sequences, as this scenario most closely resembled clinical application. As mentioned in Supplementary section 1.2.2, not all target sequences are appropriate candidates to leave out. In the case of GUIDE-seq data, one sequence fit our description: *GGTGAGT-GAGTGTGTGCGTGNGG*.

We trained logistic regression, SVM, and DNN models on the remaining data and calculated performance metrics for the left-out sequence’s data. We chose to compare only to classical machine learning baseline models because the previous state of the art models were trained on our testing sequences. Thus, comparing to these models would not be appropriate.

We noticed that the difference between the mean of the anchor-to-positive-example distances and the mean of the anchor-to-negative-example distances for the base mining model (Figure S4b) looked very similar, if not better than, that of the smart mining model (Figure S4d).

The average AUC-ROC value was 0.91, with a standard deviation of 0.025, for the base triplet loss model; the average AUC-ROC value was 0.88, with a standard deviation of 0.031, for the smart mining model (Table S1). When testing on unseen sgRNAs, the base triplet loss model had a slightly higher AUC-ROC than did the triplet loss model that employed smart mining (Figure 3b).

When comparing with baseline models, we found that crispr2vec without smart mining had the 2nd highest AUC (Figure 3b).

### 5.3 Cross assay generalization: CIRCLE-seq

We wanted to examine how well the embeddings learned by crispr2vec trained on one off-target detection assay translated to another. We split GUIDE-seq and CIRCLE-seq data into their respective symmetric differences. We trained one triplet loss model on GUIDE-seq off-target sequences that are not contained in CIRCLE-seq data. We trained another triplet loss model on CIRCLE-seq off-target sequences that are not contained in GUIDE-seq data. We tested the models on the off-target sequences contained in both GUIDE-seq and CIRCLE-seq. We visualized learned embeddings of training and testing set data for target sequences across the different models (Figure S5 depicts visualizations for one particular sequence).

For each model, we calculated the distances between pairs of cut sequences in the testing set. We correlated the distances learned by the model trained on GUIDE-seq to the distances learned by the model trained on CIRCLE-seq data. The Spearman coefficient between the learned distances was 0.60, the Pearson coefficient was 0.55, and the Kendall tau was 0.42 (Figure 5a). The distribution of distances learned by both models resemble each other (Figure 5b).

**Figure 5:**
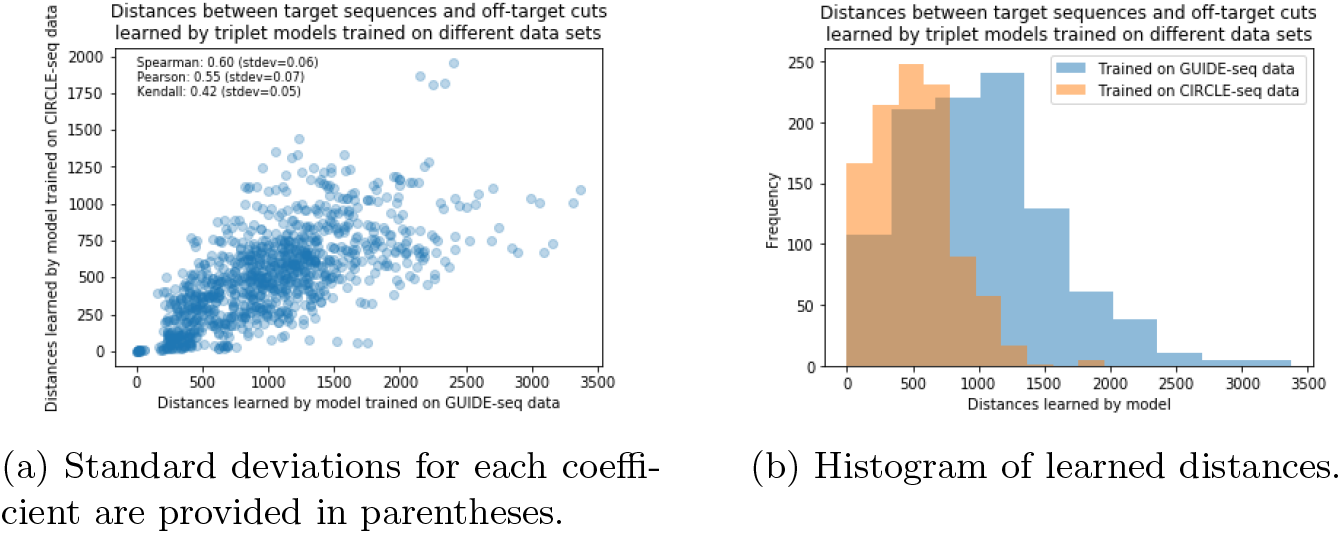
Distances between the embeddings of target sequences and off-target cuts learned by a base triplet loss model trained on GUIDE-seq data compared to those learned by a base triplet loss model trained on CIRCLE-seq data.

## 6 Discussion

crispr2vec outperforms both prior art and traditional machine learning methods, with high AUC-ROC, in the seen sgRNA context. Smart mining improves the model further, increasing the AUC from 0.96 to 0.99 (Figure 3a). Smart mining appeared to drive the positive examples closer to the anchor and the negative examples further away from the anchor (Figure 4d). The reported results for prior art can be considered an upper bound for their performance, as these models were trained on the testing data.

In the unseen sgRNA context, crispr2vec outperforms all traditional machine learning methods. Smart mining did not improve performance as it did in the seen guide RNA case. We postulate that this is because of patterns unique to the left-out target sequence’s data that are not captured by the training data. Smart mining accentuates the patterns unique to the left-out target sequence’s data that is not captured by the training data. This a statistical artifact resulting from the fact that the total distribution of mismatches in GUIDE-seq data is quite different than the distribution of mismatches in GUIDE-seq data for just the unseen sequence (Figure S2c). A big limitation here is the limited amount of training data available.

The unseen sgRNA scenario warrants further explanation. crispr2vec out-performed all baseline models except Hamming distance (Figure 3b). While Hamming distance is an interesting theoretical baseline, it’s not particularly useful in practice. Hamming distance classifies too many true non-cuts as cuts; its specificity is too high.

In all cases, crispr2vec outperformed the traditional neural network baseline. The DNN was a multi-layer dense neural network with the same architecture as crispr2vec; but it takes as input a single off-target sequence and outputs a classification as cut/not-cut. This reinforced our choice of a metric learning approach, since incorporating the intended target into our training enhanced the performance of the neural network.

Our model was, indeed, flexible enough to capture the inherent cutting dynamics of the sequences. The distribution of distances learned by crispr2vec on GUIDE-seq data closely resembled the distribution of distances learned by crispr2vec on CIRCLE-seq data (Figure 5b). The Spearman correlation for the distances learned by crispr2vec was comfortably above 0.5 (Figure 5a). We are learning comparable sequence distances and embeddings between the two assays. Thus, the distances learned by our model may be able to be agnostic to the assay from which the data originates.

Further, any linear transform from a pair of one-hot encoded vectors can be thought of as a weighted inhomogenous Hamming distance between two sequences. In this case, since we use ReLUs instead of linear transforms, we have essentially learned a weighted inhomogenous Hamming distance between sequences. To the best of our knowledge, such a model has not been used elsewhere.

Overall, crispr2vec achieved an average AUC-ROC value of 0.99 when testing on seen target sequences and 0.91 when testing on unseen target sequences. The latter results, in particular, exceed the reported state-of-the-art in CRISPR off-target modeling. The learned embeddings generalize across other detection assays and may be useful for other tasks, such as the prediction of CRISPR sgRNA sensitivity to a target site.

## 7 Conclusion

We presented **crispr2vec**, which CRISPR-like sequences. We employed a metric learning approach, for which we introduced a offline, smart triplet sampling strategy. We demonstrated superiority over previous state-of-the-art methods in predicting CRISPR off-target effects. Finally, we showed the potential for our learned embeddings to generalize across off-target detection assays.

This work lends itself naturally to a number of future investigations. We posit that the integration of additional features, such as location and epigenetics, would augment the embeddings learned by crispr2vec. We plan to expand our model to incorporate convolutions and sequence-based modeling to improve performance and generalization further. The datasets could be expanded to more assays, more model organisms, and other CRISPR-Cas systems to further explore whether a trained triplet loss model is generalizable. Additionally, applying new machine learning interpretability tools to our model may further illuminate the underlying biology of the cutting dynamics of CRISPR-Cas systems captured by crispr2vec. In addition, we could use the learned embeddings to predict CRISPR sensitivity for guide-target affinities.

Time and again, interdisciplinary areas have moved forward with the incorporation of work in other applied fields. We hope that this work encourages more such creativity, especially as the rise of CRISPR-based therapeutics makes the need for predicting cleavage patterns more immediate and urgent.

## References

[1] Shiran Abadi, Winston X. Yan, David Amar, and Itay Mayrose. A machine learning approach for predicting crispr-cas9 cleavage efficiencies and patterns underlying its mechanism of action. PLoS computational biology, 13, 2017.

[2] Amir Alavi, Matthew Ruffalo, Aiyappa Parvangada, Zhilin Huang, and Ziv Bar-Joseph. A web server for comparative analysis of single-cell rna-seq data. Nature communications, 9(1):1–11, 2018.

[3] Babak Alipanahi, Andrew Delong, Matthew T Weirauch, and Brendan J Frey. Predicting the sequence specificities of dna-and rna-binding proteins by deep learning. Nature biotechnology, 33(8):831–838, 2015.

[4] Sangsu Bae, Jeongbin Park, and Jin-Soo Kim. Cas-offinder: a fast and versatile algorithm that searches for potential off-target sites of cas9 rna-guided endonucleases. Bioinformatics, 30(10):1473–1475, 2014.

[5] James P Broughton, Xianding Deng, Guixia Yu, Clare L Fasching, Venice Servellita, Jasmeet Singh, Xin Miao, Jessica A Streithorst, Andrea Grana-dos, Alicia Sotomayor-Gonzalez, et al. CRISPR–Cas12-based detection of SARS-CoV-2. Nature Biotechnology, pages 1–5, 2020.

[6] Janice S Chen, Yavuz S Dagdas, Benjamin P Kleinstiver, Moira M Welch, Alexander A Sousa, Lucas B Harrington, Samuel H Sternberg, J Keith Joung, Ahmet Yildiz, and Jennifer A Doudna. Enhanced proofreading governs crispr–cas9 targeting accuracy. Nature, 550(7676):407–410, 2017.

[7] Sidi Chen, Neville E Sanjana, Kaijie Zheng, Ophir Shalem, Kyungheon Lee, Xi Shi, David A Scott, Jun Song, Jen Q Pan, Ralph Weissleder, et al. Genome-wide crispr screen in a mouse model of tumor growth and metastasis. Cell, 160(6):1246–1260, 2015.

[8] Weihua Chen, Xiaotang Chen, Jianguo Zhang, and Kaiqi Huang. Beyond triplet loss: a deep quadruplet network for person re-identification. In Proceedings of the IEEE Conference on Computer Vision and Pattern Recognition, pages 403–412, 2017.

[9] De Cheng, Yihong Gong, Sanping Zhou, Jinjun Wang, and Nanning Zheng. Person re-identification by multi-channel parts-based cnn with improved triplet loss function. In Proceedings of the iEEE conference on computer vision and pattern recognition, pages 1335–1344, 2016.

[10] Seung Woo Cho, Sojung Kim, Yongsub Kim, Jiyeon Kweon, Heon Seok Kim, Sangsu Bae, and Jin-Soo Kim. Analysis of off-target effects of crispr/cas-derived rna-guided endonucleases and nickases. Genome research, 24(1):132–141, 2014.

[11] Le Cong, F Ann Ran, David Cox, Shuailiang Lin, Robert Barretto, Naomi Habib, Patrick D Hsu, Xuebing Wu, Wenyan Jiang, Luciano A Marraffini, et al. Multiplex genome engineering using crispr/cas systems. Science, 339(6121):819–823, 2013.

[12] Jacob Devlin, Ming-Wei Chang, Kenton Lee, and Kristina Toutanova. Bert: Pre-training of deep bidirectional transformers for language understanding. arXiv preprint arXiv:1810.04805, 2018.

[13] Prafulla Dhariwal, Heewoo Jun, Christine Payne, Jong Wook Kim, Alec Radford, and Ilya Sutskever. Jukebox: A generative model for music. arXiv preprint arXiv:2005.00341, 2020.

[14] Atray Dixit, Oren Parnas, Biyu Li, Jenny Chen, Charles P Fulco, Livnat Jerby-Arnon, Nemanja D Marjanovic, Danielle Dionne, Tyler Burks, Raktima Raychowdhury, et al. Perturb-seq: dissecting molecular circuits with scalable single-cell rna profiling of pooled genetic screens. Cell, 167(7):1853–1866, 2016.

[15] John G Doench, Nicolo Fusi, Meagan Sullender, Mudra Hegde, Emma W Vaimberg, Katherine F Donovan, Ian Smith, Zuzana Tothova, Craig Wilen, Robert Orchard, et al. Optimized sgrna design to maximize activity and minimize off-target effects of crispr-cas9. Nature biotechnology, 34(2):184, 2016.

[16] Jennifer A Doudna and Emmanuelle Charpentier. The new frontier of genome engineering with crispr-cas9. Science, 346(6213):1258096, 2014.

[17] Gökcen Eraslan, Žiga Avsec, Julien Gagneur, and Fabian J Theis. Deep learning: new computational modelling techniques for genomics. Nature Reviews Genetics, 20(7):389–403, 2019.

[18] Richard L Frock, Jiazhi Hu, Robin M Meyers, Yu-Jui Ho, Erina Kii, and Frederick W Alt. Genome-wide detection of dna double-stranded breaks induced by engineered nucleases. Nature biotechnology, 33(2):179, 2015.

[19] Chuai G, Ma H, Yan J, Chen M, Hong N, Xue D, Zhou C, Zhu C, Chen K, Duan B, Gu F, Qu S, Huang D, Wei J, and Liu Q. DeepCRISPR: optimized CRISPR guide RNA design by deep learning. Genome biology, 19(1):80, 2018.

[20] Caixia Gao. The future of crispr technologies in agriculture. Nat Rev Mol Cell Biol, 19(5):275–276, 2018.

[21] Maximilian Haeussler, Kai Schönig, Héelène Eckert, Alexis Eschstruth, Joffrey Mianné, Jean-Baptiste Renaud, Sylvie Schneider-Maunoury, Alena Shkumatava, Lydia Teboul, Jim Kent, et al. Evaluation of off-target and on-target scoring algorithms and integration into the guide rna selection tool crispor. Genome biology, 17(1):148, 2016.

[22] Patrick D Hsu, Eric S Lander, and Feng Zhang. Development and applications of crispr-cas9 for genome engineering. Cell, 157(6):1262–1278, 2014.

[23] Patrick D Hsu, David A Scott, Joshua A Weinstein, F Ann Ran, Silvana Konermann, Vineeta Agarwala, Yinqing Li, Eli J Fine, Xuebing Wu, Ophir Shalem, et al. Dna targeting specificity of rna-guided cas9 nucleases. Nature biotechnology, 31(9):827, 2013.

[24] Martin Jinek, Krzysztof Chylinski, Ines Fonfara, Michael Hauer, Jennifer A Doudna, and Emmanuelle Charpentier. A programmable dual-rna–guided dna endonuclease in adaptive bacterial immunity. science, 337(6096):816–821, 2012.

[25] Julia Joung, Alim Ladha, Makoto Saito, Michael Segel, Robert Bruneau, W Huang Meei-li, Nam-Gyun Kim, Xu Yu, Jonathan Li, Bruce D Walker, et al. Point-of-care testing for covid-19 using sherlock diagnostics. medRxiv, 2020.

[26] David R Kelley, Jasper Snoek, and John L Rinn. Basset: learning the regulatory code of the accessible genome with deep convolutional neural networks. Genome research, 26(7):990–999, 2016.

[27] Daesik Kim, Sojung Kim, Sunghyun Kim, Jeongbin Park, and Jin-Soo Kim. Genome-wide target specificities of crispr-cas9 nucleases revealed by multiplex digenome-seq. Genome research, 26(3):406–415, 2016.

[28] Hui Kwon Kim, Younggwang Kim, Sungtae Lee, Seonwoo Min, Jung Yoon Bae, Jae Woo Choi, Jinman Park, Dongmin Jung, Sungroh Yoon, and Hyongbum Henry Kim. SpCas9 activity prediction by DeepSpCas9, a deep learning-based model with high generalization performance. Science Advances, 5(11), 2019.

[29] Benjamin P Kleinstiver, Vikram Pattanayak, Michelle S Prew, Shengdar Q Tsai, Nhu T Nguyen, Zongli Zheng, and J Keith Joung. High-fidelity crispr–cas9 nucleases with no detectable genome-wide off-target effects. Nature, 529(7587):490–495, 2016.

[30] Gavin J Knott and Jennifer A Doudna. Crispr-cas guides the future of genetic engineering. Science, 361(6405):866–869, 2018.

[31] Alex Krizhevsky, Ilya Sutskever, and Geoffrey E Hinton. Imagenet classification with deep convolutional neural networks. In Advances in neural information processing systems, pages 1097–1105, 2012.

[32] Jennifer Listgarten, Michael Weinstein, Benjamin P Kleinstiver, Alexander A Sousa, J Keith Joung, Jake Crawford, Kevin Gao, Luong Hoang, Melih Elibol, John G Doench, et al. Prediction of off-target activities for the end-to-end design of CRISPR guide RNAs. Nature biomedical engineering, 2(1):38–47, 2018.

[33] Laurens van der Maaten and Geoffrey Hinton. Visualizing data using t-sne. Journal of machine learning research, 9(Nov):2579–2605, 2008.

[34] Prashant Mali, Luhan Yang, Kevin M Esvelt, John Aach, Marc Guell, James E DiCarlo, Julie E Norville, and George M Church. Rna-guided human genome engineering via cas9. Science, 339(6121):823–826, 2013.

[35] Tessa G Montague, José M Cruz, James A Gagnon, George M Church, and Eivind Valen. Chopchop: a crispr/cas9 and talen web tool for genome editing. Nucleic acids research, 42(W1):W401–W407, 2014.

[36] Miguel A. Moreno-Mateos, Charles E. Vejnar, Jean-Denis Beaudoin, Juan P. Fernandez, Emily K. Mis, Mustafa K. Khokha, and Antonio J. Giraldez. CRISPRscan: designing highly efficient sgRNAs for CRISPR/Cas9 targeting in vivo. Nature methods, 12(10):982–988, 2015.

[37] Aaron van den Oord, Sander Dieleman, Heiga Zen, Karen Simonyan, Oriol Vinyals, Alex Graves, Nal Kalchbrenner, Andrew Senior, and Koray Kavukcuoglu. Wavenet: A generative model for raw audio. arXiv preprint arXiv:1609.03499, 2016.

[38] Caroline F Peddle and Robert E MacLaren. Focus: genome editing: the application of crispr/cas9 for the treatment of retinal diseases. The Yale journal of biology and medicine, 90(4):533, 2017.

[39] Ryan Poplin, Pi-Chuan Chang, David Alexander, Scott Schwartz, Thomas Colthurst, Alexander Ku, Dan Newburger, Jojo Dijamco, Nam Nguyen, Pegah T Afshar, et al. A universal snp and small-indel variant caller using deep neural networks. Nature biotechnology, 36(10):983–987, 2018.

[40] F Ann Ran, Le Cong, Winston X Yan, David A Scott, Jonathan S Gootenberg, Andrea J Kriz, Bernd Zetsche, Ophir Shalem, Xuebing Wu, Kira S Makarova, et al. In vivo genome editing using staphylococcus aureus cas9. Nature, 520(7546):186–191, 2015.

[41] F Ann Ran, Patrick D Hsu, Jason Wright, Vineeta Agarwala, David A Scott, and Feng Zhang. Genome engineering using the crispr-cas9 system. Nature protocols, 8(11):2281, 2013.

[42] Florian Schroff, Dmitry Kalenichenko, and James Philbin. Facenet: A unified embedding for face recognition and clustering. In Proceedings of the IEEE conference on computer vision and pattern recognition, pages 815–823, 2015.

[43] Gerald Schwank, Bon-Kyoung Koo, Valentina Sasselli, Johanna F Dekkers, Inha Heo, Turan Demircan, Nobuo Sasaki, Sander Boymans, Edwin Cuppen, Cornelis K van der Ent, et al. Functional repair of cftr by crispr/cas9 in intestinal stem cell organoids of cystic fibrosis patients. Cell stem cell, 13(6):653–658, 2013.

[44] Ritambhara Singh, Jack Lanchantin, Gabriel Robins, and Yanjun Qi. Deepchrome: deep-learning for predicting gene expression from histone modifications. Bioinformatics, 32(17):i639–i648, 2016.

[45] Ian M Slaymaker, Linyi Gao, Bernd Zetsche, David A Scott, Winston X Yan, and Feng Zhang. Rationally engineered cas9 nucleases with improved specificity. Science, 351(6268):84–88, 2016.

[46] Manuel Stemmer, Thomas Thumberger, Maria del Sol Keyer, Joachim Wittbrodt, and Juan L Mateo. CCTop: an intuitive, flexible and reliable CRISPR/Cas9 target prediction tool. PloS one, 10(4), 2015.

[47] Shengdar Q Tsai, Nhu T Nguyen, Jose Malagon-Lopez, Ved V Topkar, Martin J Aryee, and J Keith Joung. Circle-seq: a highly sensitive in vitro screen for genome-wide crispr–cas9 nuclease off-targets. Nature methods, 14(6):607, 2017.

[48] Shengdar Q Tsai, Zongli Zheng, Nhu T Nguyen, Matthew Liebers, Ved V Topkar, Vishal Thapar, Nicolas Wyvekens, Cyd Khayter, A John Iafrate, Long P Le, et al. Guide-seq enables genome-wide profiling of off-target cleavage by crispr-cas nucleases. Nature biotechnology, 33(2):187, 2015.

[49] Xiaoling Wang, Yebo Wang, Xiwei Wu, Jinhui Wang, Yingjia Wang, Zhaojun Qiu, Tammy Chang, He Huang, Ren-Jang Lin, and Jiing-Kuan Yee. Unbiased detection of off-target cleavage by crispr-cas9 and talens using integrase-defective lentiviral vectors. Nature biotechnology, 33(2):175, 2015.

[50] Hao Yin, Chun-Qing Song, Joseph R Dorkin, Lihua J Zhu, Yingxiang Li, Qiongqiong Wu, Angela Park, Junghoon Yang, Sneha Suresh, Aizhan Bizhanova, et al. Therapeutic genome editing by combined viral and non-viral delivery of crispr system components in vivo. Nature biotechnology, 34(3):328–333, 2016.

[51] Haoyang Zeng, Matthew D Edwards, Ge Liu, and David K Gifford. Convolutional neural network architectures for predicting dna–protein binding. Bioinformatics, 32(12):i121–i127, 2016.

[52] Haoyang Zeng and David K Gifford. Predicting the impact of non-coding variants on dna methylation. Nucleic acids research, 45(11):e99–e99, 2017.

[53] Jian Zhou and Olga G Troyanskaya. Predicting effects of noncoding variants with deep learning–based sequence model. Nature methods, 12(10):931–934, 2015.

[54] James Zou, Mikael Huss, Abubakar Abid, Pejman Mohammadi, Ali Torkamani, and Amalio Telenti. A primer on deep learning in genomics. Nature genetics, 51(1):12–18, 2019.

