## Supplementary Information for "Crispr2vec: Machine Learning Model Predicts Off-Target Cuts of CRISPR systems"

### Supplementary Materials

#### 1 Methods

##### 1.1 Datasets

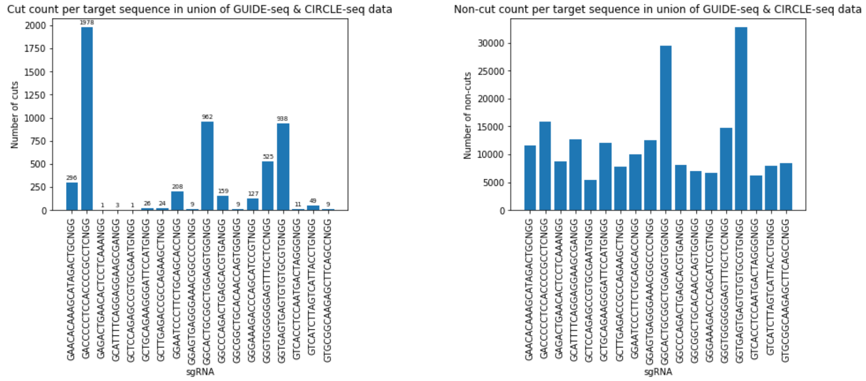


(a) Cut count per target sequence in GUIDE-seq and CIRCLE-seq

(b) Non-cut count per target sequence in GUIDE-seq and CIRCLE-seq

Figure S1: CRISPR cut sites.

##### 1.2 Model

The basic idea here is that we pass in the anchor, a positive example, and a negative example to the network, for which a distance will be calculated between samples. The loss is defined as the difference between the distance between the positive example and the anchor and the negative example and the anchor. However, optimizing just for this loss retrieves a trivial embedding that maps all points to 0. Thus, the convention is to set up the loss as the maximum of a margin (nominally) and the difference in distances.

In general for triplet loss networks, triplets from the data are passed into a neural network $f : \mathcal{R}^D \rightarrow \mathcal{R}^P$, where $D$ is the dimension of the data and $P$ is the dimension of the embedding. The loss is defined as

$$\mathcal{L}(a, p, n) = \max(d(f(a), f(p)) - d(f(a), f(n)) + \alpha, 0)$$

Where $a$ represents the anchor, $p$ represents a positive example (a sample from the same class as the anchor), and $n$ represents a negative example (a sample from a different class as the anchor). This forms the triplet $(a, p, n)$. We use our on-target sequence as the anchor, off-target sequences that are **cut** as positive examples, and off-target sequences that are **not cut** as negative examples. $\alpha > 0$ is a the margin hyperparameter. In particular, $\alpha$ is set to prevent the degenerate solution of $d(x_1, x_2) = 0$, $\forall\, x_1, x_2$. We choose $\alpha$ so that learned classes are sufficiently separated.

Therefore, this loss function takes into account both positive and negative pairs at the same time. Data from the same class should be close together in embedding space; data from different classes should be far apart. The goal of triplet loss is to create an embedding that results in two examples with the same label being close together and two samples with different labels being far apart.

##### 1.2.1 Triplet sampling

In semi-hard triplet mining, triplets are selected such that the positive example is closer to the anchor than the negative one, but the negative example is not so dissimilar from the anchor, and triplets together still register a positive loss. Specifically, $(a, p, n)$ are chosen so that $d(f(a), f(p)) < d(f(a), f(n)) < d(f(a), f(p)) + \alpha$.

We obtained our putative off-targets using Cas-OFFinder [4], which finds all biologically existing similar sequences to a target sequence. Naturally, as the number of base mismatches increase, there exist more possible permutations for a similar sequence given the OFFinder selection criteria. Indeed, we observed that a large fraction of the OFFinder sequences had 5 or 6 mismatches (Figure S2b), and the number of possible sequences monotonically increases with the number of mismatches. Compared to our positive off-target cuts, we observed that very few cuts were made when the putative off-target had more than 4 mismatches (Figure S2a). Hence, we chose to sample negative examples in accordance to the distribution of mismatches in the global positive example set. We termed this offline triplet sampling strategy **smart mining**. We posited that this would improve the adversarial robustness of our model, as distances are optimized relative to off-targets that look more "alike" to actual off-targets. In addition, we believed it, in effect, would allow us to reduce the size of our training data without sacrificing performance since detected off-targets with 5 and 6 mismatches are so infrequent relative to their combinatorial prevalence.

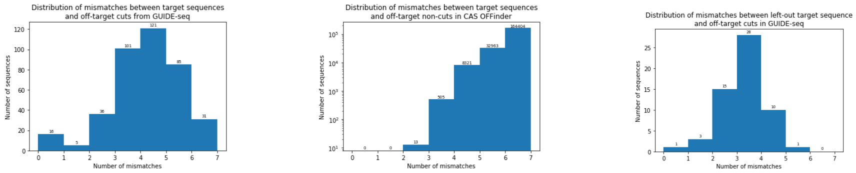


(a) Distribution over off-target cuts.

(b) Distribution over off-target non-cuts.

(c) Distribution over the unseen target sequence.

Figure S2: The distribution of mismatches between target sequences and off-target cuts in GUIDE-seq data and associated Cas-OFFinder data.

##### 1.2.2 Cross Validation Schemes

We tested the performance of our models with k-fold cross validation on seen target sequences as well as leave-one-out cross validation on unseen target sequences.

We chose $k = 5$ for cross validation on seen sequences. For each target sequence, we split the positive examples into 5 folds and the negative examples into 5 folds. We held out one positive fold and one negative fold per target sequence; we trained on the rest of the folds and tested on the held-out folds, calculating the model's accuracy on the held-out folds. This was repeated 4 more times, holding out the other positive and negative folds per target sequence. We report the results averaged across 5 splits in our cross validation study for seen guide RNA sequences. In addition, we experimented with different margin hyperparameter $\alpha$ values.

Under the leave-one-out cross validation scheme, we held out all positive and negative examples for one target sequence. We trained on the rest of the data and tested on the held-out target sequence's data. The choice of left-out target sequences was important — leave out a target with too many associated examples and the model wouldn't have enough information on which to train, leave out a target with too few associated examples and we would risk overfitting the model on the training data. For this reason, we chose to leave out target sequences associated with between 10-20% of the total number of positive examples.

##### 1.2.3 Baseline Comparisons

We compared our crispr2vec models to previous methods including CCTOP [46], CFD [15], CRISTA [1], Hsu-Zhang [23], as well as other classical machine learning methods: including logistic regression, SVM, and a regular dense neural network (DNN) with the same layer architecture as **crispr2vec** but with a binary cross entropy loss. We also considered Principle Component Analysis (PCA) as a simple embedding mechanism, where we consider the distances in the learned latent space. In addition, we considered Hamming distance between target and off-target pairs as a simple metric. We compared crispr2vec with and without smart mining to the aforementioned baseline models. We do not compare to methods such as deepCRISPR [19], DeepSpCas9 [28], and Elevation [32]. We note that while these methods would theoretically serve as good benchmarks for our method, they both use different data sources (for instance, integration of epigenetics) and the trained models these papers provide online are trained on guide-RNAs contained in the test set for our method in each scenario. Thus, there is no straightforward and fair way to provide a comparison between our method and theirs.

For seen guide RNAs, we used data splits analogous to a k-fold cross validation. We downloaded the trained models for CCTOP, CFD, CRISTA, and Hsu-Zhang from their associated papers, and applied them directly on the left out fold. Namely, we used all data for the assays in consideration when calculating performance metrics for CCTOP, CFD, CRISTA, Hsu-Zhang, Hamming distance, and PCA. We trained logistic regression, SVM, and DNN on the same data as crispr2vec, considering the off-target cut and off-target non-cut pairs as binary labels. The crispr2vec models used here were the same k-fold cross validated models as described above.

For unseen guide RNAs, we held out at each step data associated with one target sequence. We trained logistic regression, SVM, and DNN models on the remaining data and calculated performance metrics for the left-out sequence's data. We chose to compare only to classical machine learning baseline models because the previous state of the art models were trained on our testing sequences. We reported the average performance across all possible left-out sequences. The crispr2vec models used here were the same leave-one-out cross validated models as described above.

### 2 Results

| **Method** | **Data Set** | **# Repeats** | **Smart Mining** | **Margin** | **AUC (stdev)** | **Precision (stdev)** | **Recall (stdev)** | **F1 (stdev)** |
|---|---|---|---|---|---|---|---|---|
| k-fold | GUIDE-seq | 5 | False | 1.0 | 0.96 (0.012) | 1.0 (0.0) | 0.98 (0.0083) | 0.99 (0.0042) |
| | | 5 | True | 1.0 | 0.99 (0.0041) | 1.0 (0.0) | 0.99 (0.0062) | 0.99 (0.0032) |
| | GUIDE-seq & CIRCLE-seq | 5 | False | 1.0 | 0.94 (0.013) | 1.0 (0.0) | 0.97 (0.0093) | 0.98 (0.0049) |
| | | 5 | True | 1.0 | 0.99 (0.0021) | 1.0 (0.0) | 0.99 (0.0021) | 0.99 (0.0011) |
| LOOCV | GUIDE-seq | 10 | False | 1.0 | 0.91 (0.025) | 1.0 (0.0) | 0.92 (0.024) | 0.96 (0.013) |
| | | 10 | True | 1.0 | 0.89 (0.031) | 1.0 (0.0) | 0.88 (0.032) | 0.94 (0.019) |
| | GUIDE-seq & CIRCLE-seq | 10 | False | 1.0 | 0.79 (0.012) | 1.0 (0.0) | 0.79 (0.012) | 0.88 (0.0074) |
| | | 10 | True | 1.0 | 0.77 (0.044) | 1.0 (0.0) | 0.77 (0.044) | 0.87 (0.030) |

Table S1: Summary of crispr2vec methods.

#### 2.1 Hyperparameter Tuning

When tuning hyperparameters of the model, we focused on the loss function's margin $\alpha$ value. The resulting AUC-ROC values were approximately the same across all values of $\alpha$ (that ranged from 0.3 to 50). This was true for both the base triplet loss model (Figure S3a) as well as for the triplet loss model with smart mining (Figure S3b). As such, we henceforth used a margin $\alpha$ value of 1.0.

In addition, we experimented with different dropout rates. We chose a base triplet loss model with no dropout regularization steps and a smart mining triplet loss model with two dropout regularization steps (each with a 20% dropout rate).

#### 2.2 Testing on Unseen sgRNAs

See Figure S4 for additional figures.

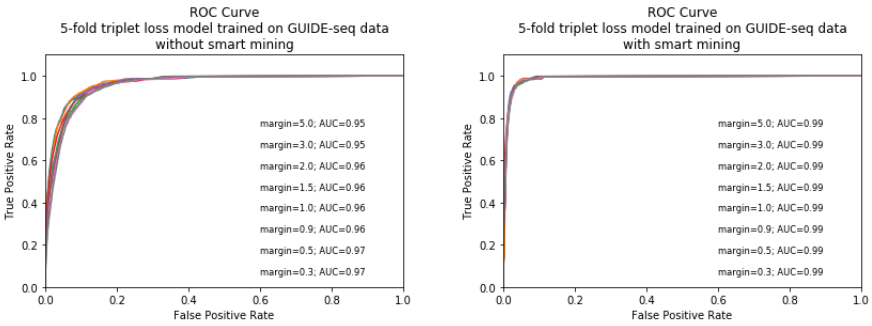


(a) Model trained without smart triplet mining

(b) Model trained using smart triplet mining

Figure S3: ROC curves for 5-fold triplet loss models on GUIDE-seq data with different margin $\alpha$ hyperparameter values.

#### 2.3 Cross assay generalization: CIRCLE-seq

See Figure S5 for additional figures.

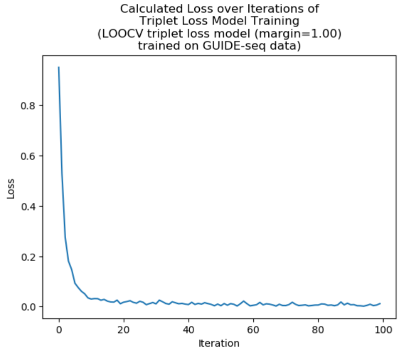


(a) Calculated loss over iterations of the triplet loss model training phase.

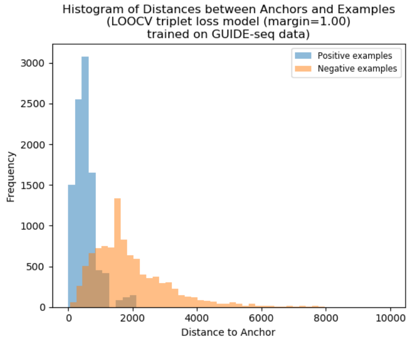


(b) Histogram of the calculated distances between anchors to positive examples and anchors to negative examples in the testing set.

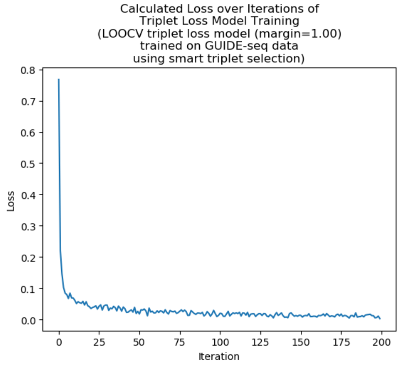


(c) Calculated loss over iterations of the triplet loss model smart mining training phase.

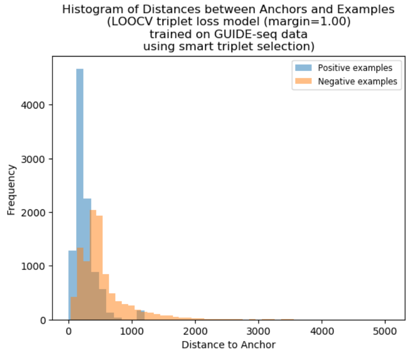


(d) Histogram of the calculated distances between anchors to positive examples and anchors to negative examples in the testing set when trained with smart mining.

Figure S4: Leave-one-out cross validation of crispr2vec trained on GUIDE-seq data.

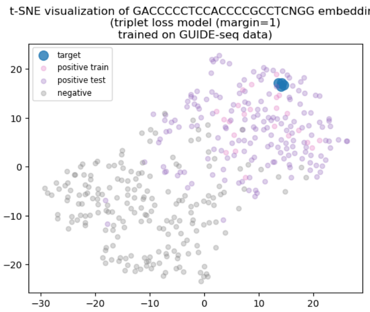


(a) crispr2vec trained on GUIDE-seq data.

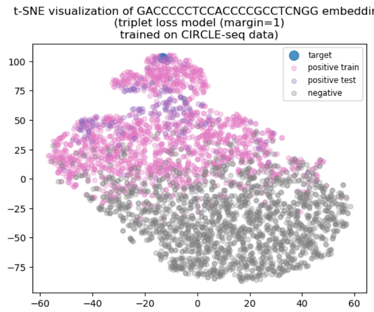


(b) crispr2vec trained on CIRCLE-seq data.

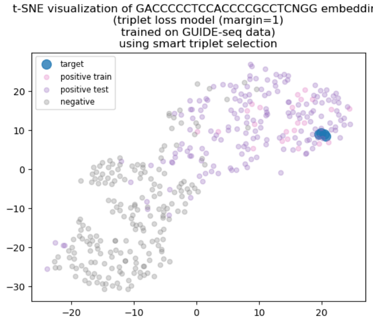


(c) crispr2vec trained on GUIDE-seq data with smart mining.

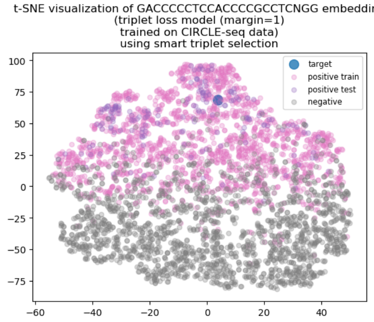


(d) crispr2vec trained on CIRCLE-seq data with smart mining.

Figure S5: t-SNE visualizations of a target sequence.
